# Evolution of bone cortical compactness in slow arboreal mammals

**DOI:** 10.1101/2020.09.02.279836

**Authors:** F. Alfieri, J.A. Nyakatura, E. Amson

**Author notes:** Corresponding author: Fabio Alfieri.

## Abstract

Lifestyle convergences and related phenotypes are a major topic in evolutionary biology. Low bone cortical compactness (CC), shared by the two genera of ‘tree sloths’, has been linked to their convergently evolved slow arboreal ecology. The proposed relationship of low CC with ‘tree sloths’ lifestyle suggests a potential convergent acquisition of this trait in other slow arboreal mammals. ‘Tree sloths’, ‘Lorisidae’, Palaeopropithecidae, *Megaladapis* and koalas were included in a phylogenetically informed CC analysis of the humerus and femur, as well as closely related but ecologically distinct taxa. We found that ‘tree sloths’ are unparalleled by any analysed clade, in featuring an extremely low CC. A tendency for low CC was however found in palaeopropithecids (especially *Palaeopropithecus*) and *Megaladapis*. On the other hand, low CC was not found in ‘Lorisidae’. Koalas, although deviating from the compact structure generally observed in mammals, are not clearly distinct from their sister taxon (wombats) and show humeral CC that is higher than femoral CC. Multiple factors seem to influence CC, preventing the recognition of a simple relationship between slow arborealism and low CC. Highly compact cortical bone in extinct sloths confirms that low CC in ‘tree sloths’ likely represents a recent convergence.

## 1. Background

Tetrapod long bones microstructure is known to reflect life history, physiology, phylogeny, ontogenetic growth and lifestyle [1–3]. Indeed, diaphyseal microstructure has been related to ecology [2,4,5] and employed to infer it in extinct tetrapods (e.g. [6,7]). Moreover, this relationship potentially enables to recognise patterns of convergent evolution [8].

To highlight convergent phenotypes, mammals are particularly suitable to be investigated given their remarkable diversity of lifestyles. Mammalian groups with different ecological adaptations can be discriminated analyzing several diaphyseal microstructural parameters (e.g. [9,10]). The amount of porosities in the diaphyseal compact cortical bone, or cortical compactness (CC), was recently quantified in the humerus of adult extant and extinct xenarthrans. The two lineages of extant sloths (*Choloepus* and *Bradypus*), characterised by an exceptionally slow arboreal lifestyle [11], were found to feature lower CC in respect to other similar-sized taxa which conversely follow the generalised mammalian condition of high CC [12]. Because ‘tree sloths’ are not monophyletic (hence the quotation marks employed herein), this trait was identified as one of the convergent traits related to their independently acquired slow arboreal lifestyle [11,12]. If indeed driven by adaptation to this lifestyle, a low CC may have convergently evolved in other slow arboreal therians, too. With ‘Slow Arboreal’, we refer to species characterised by exceptionally slow and cautious movements on trees and daily activities budgets dominated by rest and quiescence. This set of traits is generally associated with unusually low metabolic rate. To date, no CC analyses were performed on slow arboreal mammals, beside the one on ‘tree sloths’ [12]. Following the latter, we quantified CC in the humerus. Moreover, we also investigated CC of the femur, since this bone is often employed in microstructural analysis (e.g. [13,14]).

Together with ‘tree sloths’, we analysed other known slow arboreal lineages, namely the koala *Phascolarctos cinereus* [15] and ‘lorisids’ [16], with the latter arguably representing a paraphyletic group (see Materials and Methods). Moreover, taxa from two late Quaternary-Holocene subfossil lineages of Malagasy strepsirrhines, Palaeopropithecidae (*Palaeopropithecus, Mesopropithecus* and *Babakotia*) and Megaladapidae (*Megaladapis*) [17,18], are here investigated. They were described as slow and poorly agile arboreal, through postcranial (gross anatomy, e.g. [17,19], diaphyseal and epiphyseal geometric properties, [20]) and cranial [21] evidences (reviewed in [18,20]). Beside the taxa introduced above, a set of close relatives, conversely characterised by different lifestyles, is also studied (Figure 1, Table S1). Taking into account phylogenetic relationships, this ecologically heterogeneous sample enables to identify convergent acquisitions of slow arboreality in the sampled taxa and to highlight a potential pattern of convergent low CC in slow arboreal therians. Noticeably, the sample characterised by distinct lifestyles includes specimens of extinct sloths from the Patagonian Early Miocene Santa Cruz Formation [22–24]. Analyzing them can additionally inform on the evolution of low CC in the two lineages of ‘tree sloths’ and corroborate the proposed recent convergence [12].

**Figure 1.**
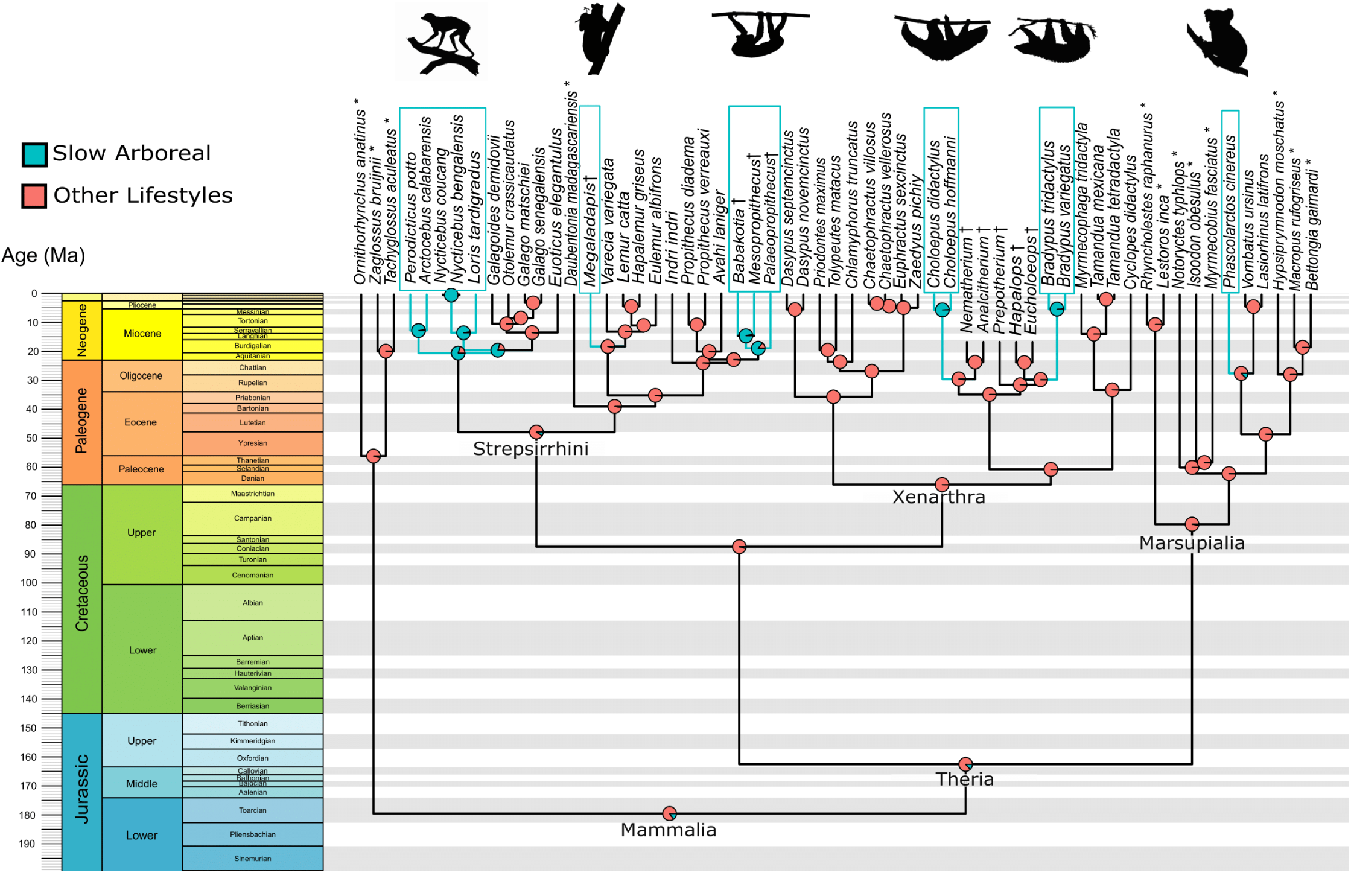
Time-calibrated tree of mammalian taxa here analysed. Six events of convergent acquisition of ‘Slow Arboreal’ adaptations were reconstructed with Stochastic Character Mapping [26]. It is shown by light blue branches and rectangles. Taxa not sampled for the cortical compactness analysis are indicated with *. The figure was built with the ‘geoscalePhylo’ function (‘strap’ R 3.6.3. package, [27]) and modified in Inkscape [28].

In this study, humeral and femoral CC are quantified in a wide mammalian sample, representing slow arboreal clades and near relatives with diverging lifestyles. Results are interpreted in a phylogenetically informed manner. In this way, slow arboreality convergences could be reconstructed and ecological effects on CC are preponderantly taken into account. If a direct relationship with this extreme lifestyle is present, we expect to find the convergent slow arboreal groups consistently showing lower CC in respect to close relatives.

## 2. Material and Methods

### a. Sampled collections

Ten mammals collections were sampled: Museum für Naturkunde, Berlin (ZMB), Staatliches Museum für Naturkunde, Stuttgart (SMNS), Zoologisches Forschungsmuseum Alexander Koenig, Bonn (ZFMK), Zoologische Staatssammlung, Munich (ZSM), Germany; Naturhistorisches Museum, Wien (NMW), Austria; Muséum national d’Histoire naturelle, Paris (MNHN), France; American Museum of Natural History, New York, NY, (AMNH), Field Museum of Natural History, Chicago, IL, (FMNH), Yale Peabody Museum of Natural History, New Haven, CT, (YPM-PU) and Division of Fossil Primates, Duke Lemur Center, Durham. NC (DPC), USA. A total of 106 humeri and 106 femora, belonging to 47 taxa were sampled. Only non-pathological, non-captive and adult individuals were included. Adulthood was established assessing the degree of epiphyseal fusion, in general, and bone size for marsupials, given their incomplete epiphyseal fusion through adulthood [25].

### b. Data acquisition

Bones were scanned using micro Computed-Tomography (µCT) (Phoenix | x-ray Nanotom, GE Sensing and Inspection Technologies GmbH; XYLON FF35-CT-System, YXLON GmbH; Microtomograph RX EasyTom 150; Nikon XTH 225 ST; GE v|tome|x). CC being acquired at mid-diaphysis (see below), the field of CT-scan acquisition was focused on this region, in order to reach the highest possible resolution. The mid-shaft was defined as 50% of the whole bone length (Figure S1). Image stacks (16-bit tifs) were acquired with a resolution ranging from 0.0045 to 0.0662 mm (Table S2 and S3), from which the 50% slice was taken. This resolution range typically allows capturing all vacuities except for the smaller lumen of osteons [12] and it is here confirmed for lower resolutions as well. The resolution is unavoidably affected by interspecific differences in bone gross anatomy. Indeed, the presence of bony processes at mid-diaphysis (e.g. third trochanter in the femur of Cingulata) forces an acquisition with longer specimen to X-ray source distance, thus with lower resolution. For one specimen (femur of *Priodontes maximus* ZSM 1931-293) the resolution was purposely increased excluding the mid-diaphysis process from the acquired field of view (not needed to compute CC, see below and Supplementary Materials Figures S2, S3 and S4). Since its CC falls within the range of Cingulata (Table S3), we will assume that the associated bias is minor.

### c. Data processing

Following Montañez-Rivera et al. [12], all mid-shaft cross-sections were imported into Fiji [29] and automatically binarised (‘Optimize Threshold’ function, BoneJ plugin, [30]). It was effective for almost all specimens. Manual correction was necessary for two specimens. For one humerus and one femur of *Palaeopropithecus*, dense, non-osseous material was present (clearly identifiable before binarisation). The corresponding area was selected (‘Threshold’ and ‘Create Selection’ Fiji functions) and deleted (i.e. grey values set to 0), before running ‘Optimize Threshold’. Post-binarisation corrections were performed for about 15% and 25% of the humeri and femora, respectively (specimens with PostBinarCorr = ‘Yes’ in Tables S2 and S3). In such cases, since non-osseous material was recognised as ‘bone’ by the thresholding, it was manually deleted from the slices after having confirmed its nature observing images before thresholding and regions contiguous to the mid-shaft.

### d. Quantification of the Cortical Compactness (CC)

Following the criteria of Montañez-Rivera et al. [12], a Region Of Interest (ROI) was selected on each cross-section, including cortices and excluding spongiosa and medullary cavity (‘Wand (tracing)’ Fiji tool; Figure S1). In about 65% of the humeri and 60% of the femora, the cortex-spongiosa transition was not abrupt (specimens with SpongManualSel = ‘Yes’ in Tables S2 and S3), which required a manual selection [12]. The transition was unambiguously located connecting the most external holes considered as part of the medullary cavity (‘Polygon selection’ Fiji tool). Bony processes at the mid-shaft level, found in about 35% and 30% of the sampled humeri and femora, respectively (specimens with BoneProc= ‘Yes’ in Tables S2 and S3), were excluded by identifying the level of first cortical thinning and spongious bone appearance and using straight lines, perpendicular to the outer cortical surface. Cracks within the cortical region were not included in the ROIs. See Figure S2 for a demonstration of ROI selection criteria. For each ROI, the percentage of bone pixels relative to total area (ratio representing the cortical compactness, CC) and the cortical area (CA, reliable body size predictor [31]) were computed (respectively ‘Area Fraction’ and ‘Area’ routines, ‘Measure’ function of Fiji). All cross-sections with ROIs used for CC and CA computation are shown in Figures S3 and S4 while raw CC and CA values are listed in Tables S2 and S3.

### e. Time-tree for phylogenetic comparative methods

To take into account non-independence of observations due to phylogenetic relationships, a time-tree of the studied species was used to phylogenetically inform the statistical analysis (Figure 1). We used a Maximum Clade Credibility (MCC) DNA-only node-dated mammals phylogeny taken from the related posterior distribution of the work of Upham et al. [32] and representing 4098 species. In doing so, polytomies and unresolved nodes were avoided [32]. However, several adjustments to fit our sample were performed in Mesquite [33]. Two extant species (*Dasypus septemcinctus* and *Tamandua mexicana)* were added following recently published phylogenetic positioning and divergence times [34,35]. Moreover, extinct taxa were subsequently added based on previously published morphological phylogenies and/or analyses of recovered genetic information [36–38]. Santacrucian sloths’ families position in respect to *Choloepus* and *Bradypus* and all node ages of Folivora (extinct + extant sloths) were constrained on the tree provided by Delsuc et al. [36]. However, since none of the extinct sloths we studied were present in this tree, infra-familiar relationships followed the morphological phylogeny built by Varela et al. [37]. First Appearance Datum (FAD) and Last Appearance Datum (LAD) are respectively 17.5 and 16.3 Mya for the analysed extinct sloths, which all come from the Santa Cruz Formation [39]. Divergence times between *Hapalops* and *Eucholoeops* and between *Analcitherium* and *Nematherium* could only be placed in a period ranging from the previous node to the FAD. In such cases, the divergence time was arbitrarily set halfway between these two corresponding ages. *Mesopropithecus* and *Babakotia*, not present in the tree from Upham et al. [32], were added following the phylogeny of Herrera and Dávalos [38]. This phylogeny was used to set topology and date nodes for the (‘Indriidae’+Palaeopropithecidae) clade, with ‘Indriidae’ being paraphyletic [38]. Indeed, constraining ages only for palaeopropithecids would have resulted in a lack of consistency with the tree from Upham et al. [32], concerning their temporal relationships with ‘Indriidae’. It is noteworthy that the MCC tree used here involves that ‘Lorisidae’ are paraphyletic. Monophyly of the group is debated, with several phylogenies yielding disagreeing results (e.g. [40–42]).

### f. Ancestral Lifestyle Reconstruction

All statistical analyses were performed in R 3.6.3 [43]. Independent acquisitions of slow arboreal lifestyle were reconstructed with Stochastic Character Mapping (SCM; [26]). SCM was performed using the sampled taxa and an additional set of species (n=12) included to have a wider and more representative taxonomic sample of mammalian lifestyles (3 Monotremata, 8 Marsupialia and *Daubentonia madagascariensis*; Figure 1). We assigned a binary state Slow Arboreal/Non-Slow Arboreal to each terminal taxon, and applied the ‘make.simmap’ R function for SCM (Equal Rate model, 1000 simulations, ‘phytools’ package; [44]).

### g. Relationship between Cortical Compactness (CC) and lifestyle

The relationship between humeral/femoral CC and lifestyle (binary categorical variable: Non-Slow Arboreal or Slow Arboreal) was tested through Phylogenetic Generalized Least Square (PGLS) regressions. For both the humerus and the femur we performed a phylogenetic ANCOVA, controlling body mass effects adding CA as covariate (‘gls’ function; method “ML”, ‘nlme’ R package; [45]). We used mean species CC and CA values, both log-transformed. Individuals of extant species catalogued only at the generic level were here excluded and the regressions were phylogenetically informed using Pagel’s lambda (‘corPagel’ function, ‘ape’ R package, [46]). No significant relationship was found between CC and body mass (p-value_humerus_=0.45 and p-value_femur_=0.37). Humeral and femoral CC were visualised through boxplots (‘ggplot’ function, ‘ggplot2’ R package; [47]) and mean CC values mapped on the phylogeny (‘contMap’ function, ‘phytools’ R package; [44]), with graphs modified in Inkscape [28] for visualization (Figure 2). Supporting material and dataset are available from Figshare Digital Repository (https://doi.org/10.6084/m9.figshare.12896249.v4) [48].

**Figure 2.**
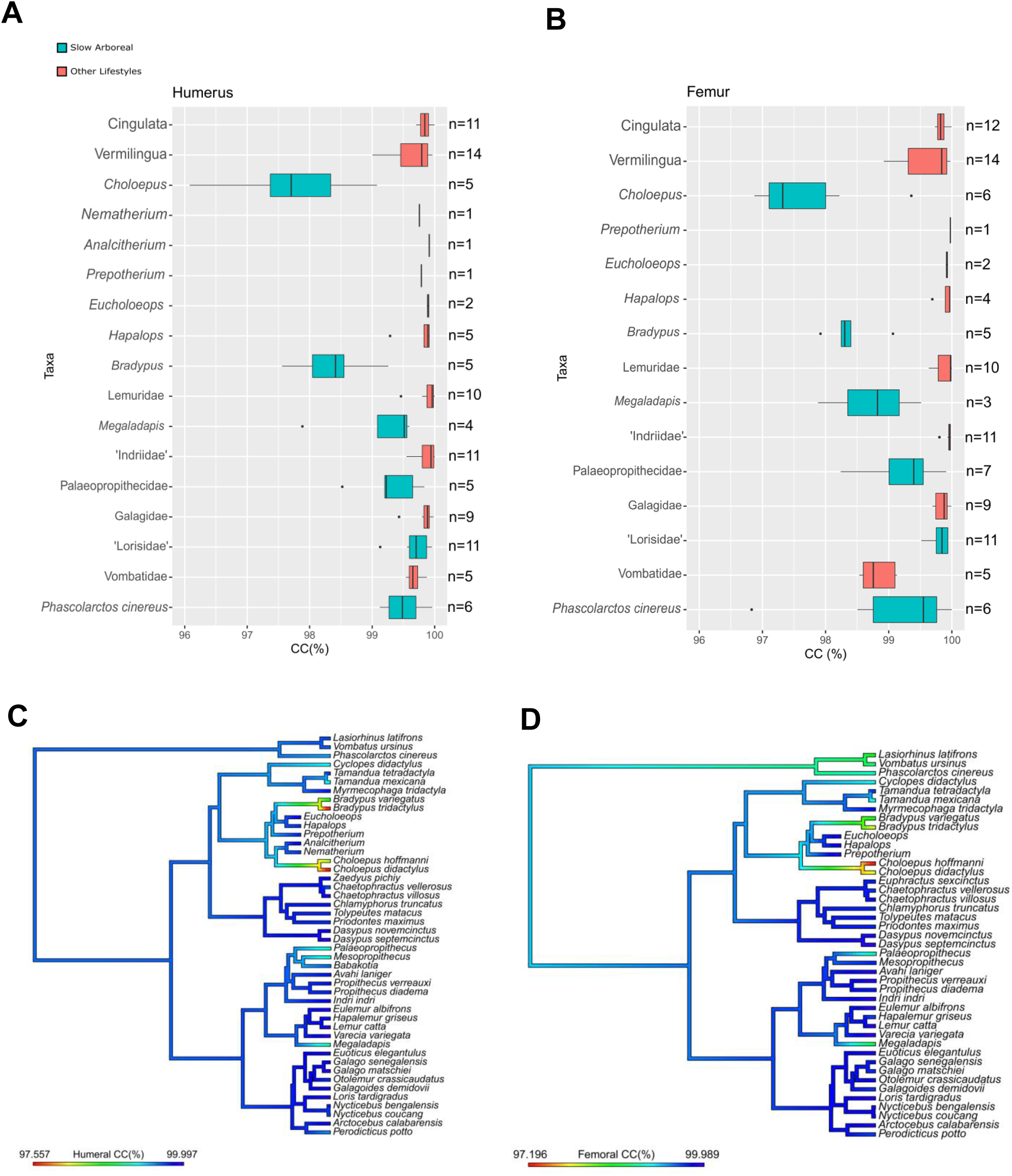
Box-and-whisker plots of humeral (A) and femoral (B) cortical compactness (CC), with specimens grouped by taxon. Black triangles and vertical bars indicate mean and median CC, respectively. B. Phylogenetic maps of the humeral (left) and femoral (right) CC averaged for each species (or genus for extinct taxa).

## 3. Results

From SCM arose a highly probable ancestral Non-Slow Arboreal condition and the convergent acquisition of slow arboreality in ‘Lorisidae’, *Megaladapis*, Palaeopropithecidae, *Choloepus, Bradypus* and *P. cinereus*, which will be referred hereafter as slow arboreal clades. The aforementioned paraphyly for ‘Lorisidae’ involves that Asian ‘Lorisidae’ (*Nycticebus* and *Loris*) are more closely related to Galagidae than to African ‘Lorisidae’ (*Arctocebus* and *Perodicticus*). Accordingly, a result of SCM is the ancestral slow arboreal condition for Lorisiformes (‘Lorisidae’ + Galagidae) with subsequent reversion in active leaping Galagidae (Figure 1).

CC values are significantly correlated with lifestyle (p-value_humerus_ = 0.4e-05; p-value_femur_ = 28e-05). However, it is obvious that not all slow arboreal clades contribute to the difference and residuals of both humeral and femoral regressions significantly deviate from normality (Shapiro-Wilk test, p-value_humerus_=2.8e-05 and p-value_femur_=50e-05).

### a. CC of ‘tree sloths’ vs. other xenarthrans

In both the humerus and the femur, ‘tree sloths’ represent the clades with the lowest CC in the entire therian sample (especially *Choloepus*, with minima of CC_humerus_ = 96.08% and CC_femur_= 96.87%; Figure 2). Their diaphyseal microstructure is characterised by numerous and large cavities, widespread in the cortex (Figures S3 and S4). ‘Tree sloths’ sharply contrast other extant (Vermilingua and Cingulata) and extinct (*Nematherium, Analcitherium, Hapalops, Eucholoeops, Prepotherium*) xenarthrans, conversely characterised by higher CC. Moreover, ‘tree sloths’ are also distinct from other xenarthrans in exhibiting a wider CC range of variability. *Choloepus* stands out on that regard as well, with a CC range of 2.99% and 2.48% for the humerus and femur, respectively (Table S1; Figure 2A, B). With high CC values and less variable CC, the other xenarthrans mirror the generalised mammalian condition. Vermilinguans also show a relatively wide range of CC variation (0.96% in the humerus and 1.05% in the femur). This is mainly due to the low values of *Cyclopes didactylus* (4 individuals with means of CC_humerus_=99.3% and CC_femur_=99.2%) and *Tamandua mexicana* (only one humerus and one femur with CC of 99.11% and 98.95%, respectively). Similarly to *Bradypus* and *Choloepus*, the vacuities of *Cy. didactylus* are widespread in the cortex and the spatial pattern is reminiscent of that of ‘tree sloths’. On the other hand, porosities of *T. mexicana* appear to be mainly concentrated toward the inner cortex, with few isolated vacuities in the outer regions (Figures S3, S4).

### b. CC of subfossil lemurs vs. other lemurs

We found low humeral and femoral mean CC in Palaeopropithecidae (especially *Palaeopropithecus*) and *Megaladapis*. It discriminates them from their closest relatives (‘Indriidae’ and Lemuridae, respectively) and other strepsirrhines who, instead, follow the generalised mammalian condition (i.e. high CC). Moreover, subfossil lemurs mirror ‘tree sloths’ also in showing a wide humeral and femoral CC variability. Although not reaching the extreme condition of *Choloepus*, both palaeopropithecids and *Megaladapis* approach the CC range of variation of *Bradypus*, with palaeopropithecids femoral CC range (1.67%) even exceeding *Bradypus* femoral CC range (1.15%; Figure 2A, B). In one femur of *Megaladapis* (MNHN MAD 1567) the repartition of the vacuities found throughout the cortex is reminiscent of that of the ‘tree sloths’ (Figure S4). Most of the other subfossil lemurs’ specimens show a heterogeneous distribution of porosities with a concentration toward the inner cortex and an almost complete absence from the periosteal region. On the other hand, *Babakotia* yielded higher CC values (data only for the humerus, 99.65%), while *Mesopropithecus* shows contrasting CC values between the humerus (lower CC, 99.2%) and the femur (higher CC, 99.77%; Figure 2C, D).

### c. CC of ‘Lorisidae’ vs. Galagidae

**‘**Lorisidae’ do not show evident differences in CC mean and variability in respect to Galagidae (Figure 2A, B). Moreover, no diverging patterns are identified by a closer inspection of different species mean CC values (Figure 2C, D).

### d. CC of koala vs. wombats

Mean CC do not clearly differ between koala (*P. cinereus*) and wombats’ (*Vombatus ursinus* and *Lasiorhinus latifrons*). However, a clear difference is observed between humeral and femoral patterns (Figure 2). Humeral mean CC is higher than femoral mean CC in both koala (CC_humerus_=99.50%, CC_femur_=99.04%) and wombats (CC_humerus_=99.68%, CC_femur_=98.82%). The femur of *P. cinereus* shows the widest range in the sample (3.17%). The humerus of the latter show a moderately wide range (0.84%), while wombats exhibit narrower CC ranges for both humerus (0.34%) and femur (0.59%). Specimens of *P. cinereus* with lower CC show a preferential distribution of the porosity toward the endosteal margin. Conversely, more porous specimens of wombats (for example, femur of *V. ursinus* SMNS 26510) show vacuities that are widespread in the whole cortex (Figures S3 and S4).

## 4. Discussion

### a. Humeral and femoral CC in ‘tree sloths’ and other xenarthrans

Our analysis of an extensive sampling of therian mammals, confirmed that ‘tree sloths’ exhibit lower CC compared to other xenarthrans, as previously shown [12], and highlights their convergent humeral and femoral CC decrease. *Choloepus*’ humerus and femur also exhibit the highest variability. None of the analysed taxa reaches ‘tree sloths’ extreme mean levels of low CC (Figure 2; Table S1). Montañez-Rivera et al. [12] related ‘tree sloths’ low CC to extensive bone remodeling, namely the continuous process involving the generation of resorption cavities (through bone resorption) and the deposition, within such spaces, of secondary osteons [49,50]. Vacuities on µCT cross-sections were identified as resorption cavities at various maturation stages recognised on histological sections [12,51]. ‘Tree sloths’ sharply contrast the general condition of terrestrial mammals in which the diaphyseal cortex shifts from highly porous and remodeled juvenile structures to lower remodeling rates in adults [12]. The latter results in few resorption cavities [49,52] and causes the generalised mammalian high CC. Conversely, low CC in adult ‘tree sloths’ suggests intense remodeling still active through adulthood.

High CC of Santacrucian extinct sloths resembles Vermilingua (but see discussion on *Cy. didactylus* and *T. mexicana* below) and Cingulata rather than more closely related ‘tree sloths’. Albeit with similar evidence, Montañez-Rivera et al. [12] did not exclude the possibility that low CC could characterise some extinct sloths. Conversely, specimens here analysed, which extend the sample of Montañez-Rivera et al. [12], coherently point toward high CC in extinct sloths. This supports the convergent acquisition of the trait in the lineages leading to *Bradypus* and *Choloepus* and it can be added to the phenotypical convergent traits shared by ‘tree sloths’ [11]. While the extinct sloths sampled here belong to three families [37], they all come from one locality (Patagonian Early Miocene Santa Cruz Formation, 17.5-16.3 Mya; [23,39]). Considering the taxonomical and ecological diversity of extinct Folivora [53], studying taxa from other ages and geographical contexts may elucidate the pattern of evolution of CC in the clade.

Low CC was found for the silky anteater, *Cy. didactylus*. Due to its elusive nocturnal habits, the ecology of *Cy. didactylus* is poorly known. However, available information suggests that it is almost fully arboreal and slow moving [54–56]. Postcranial similarities to ‘tree sloths’ have been already highlighted [57]. The convergence of low CC in ‘tree sloths’, together with slightly lower CC in *Cy. didactylus*, suggests a relationship between low CC and slow arboreality within xenarthrans. Long bone thin-sections of *Cy. didactylus* are needed to confirm that porosities observed throughout the cortex (yielding slightly lowered CC) are indeed resorption cavities or immature secondary osteons, as in ‘tree sloths’. The spatial distribution of vacuities in *Cy. didactylus* is reminiscent of the ‘tree sloths’ pattern (Figures S3 and S4). Pending histological confirmation, we tentatively propose ongoing bone remodeling in adult silky anteaters. The low CC values of *T. mexicana*, contrasting *Tamandua tetradactyla*’ s more compact structure (Figures 2C, D), is puzzling. No major ecological differences between the two (non-slow) semiarboreal *Tamandua* species are reported [58,59]. However, since CC values of *T. mexicana* come from a single individual (Table S2, S3), the possibility of biases related to small sample size should be considered.

### b. Humeral and femoral CC in slow arboreal primates and marsupials

Similarly to ‘tree sloths’, Palaeopropithecidae (mainly *Palaeopropithecus*) and *Megaladapis* show independent acquisition of humeral and femoral low CC with higher variability (Figure 2). Several postcranial convergences between subfossil lemurs and ‘tree sloths’ are documented [18–20]. Due to the abundant inferences of slow arboreal ecology for palaeopropithecids and megaladapids [17–21], low CC could be a convergence driven by their similar purported lifestyle. However, the subfossil lemurs analysed here show a CC that, despite being clearly lower than in other lemurs, is not comparable to the extreme levels of ‘tree sloths’. Only *Palaeopropithecus* and *Megaladapis* show similarly low humeral and femoral CC, while *Babakotia* and *Mesopropithecus* yielded higher CC and/or contrasting inter-limb patterns (Figure 2). One should note that *Palaeopropithecus* and *Megaladapis* are the most represented subfossil lemurs in the sample (Table S2, S3). In contrast, the femur of *Babakotia* was not analysed and the only available humerus is not well preserved. Likewise, *Mesopropithecus*’ humeri are only preserved poorly (Figure S3). Biases deriving from low number of analysed bones and specimens’ preservation should thus be taken into account. No osteo-histological analyses are available for subfossil lemurs, which prevents the characterization of the cortical vacuities responsible for low CC in *Palaeopropithecus* and *Megaladapis*. The concentration of porosities toward the inner cortex in some of their specimens matches the distribution of secondary osteons in humeral and femoral thin-sections of ‘Indriidae’ and Lemuridae [60]. This suggests that the cortical vacuities observed in subfossil lemurs are resorption cavities (immature secondary osteons). This would imply relatively intense and perennial bone remodeling being responsible for the low CC of *Palaeopropithecus* and *Megaladapis*.

Strikingly, humeral/femoral CC do not differ between ‘Lorisidae’ and Galagidae and both taxa do not deviate from the mammal general condition of highly compact/little variable adult cortex. Indeed, their highly diverging arboreal habits (slow arboreal in ‘Lorisidae’, [16] vs. active leaping in Galagidae, [61]) were considered explanatory for postcranial differences in both external and internal anatomy [62]. Moreover, several postcranial convergences between ‘Lorisidae’ and ‘tree sloths’ are reported [57,63].

Koala and wombats do not consistently differ in CC and we found differences between humeral and femoral CC, with the latter lower than the former in both koalas and wombats. Moreover, koalas exhibit the largest CC variability in our sample. Considering their ecological differences (slow arboreality of *P. cinereus* [15] vs. terrestrial fossoriality in wombats, [64]) lifestyle alone can hardly explain the CC pattern and suggest additional/alternative explanations. Differing from most marsupials, *Vombatus ursinus*’ cortex is intensely remodeled [65,66] and it is reasonable to ascribe the vacuities in wombats’ cortex to immature secondary osteons. Walker et al. [66] hypothesised that wombats’ humeral microstructure could be explained by physiological and biomechanical adaptations to fossoriality. We propose that physiological effects, systemically lowering CC through remodeling, may be locally attenuated by biomechanical needs for a compact structure in the humerus related to wombats’ forelimb based scratch-digging [66]. This scenario would be consistent with CC values lower in the femur than in the humerus, as found in *Vombatus ursinus*.

Secondary osteons are almost absent from the cortex of *P. cinereus* [65], contrasting with xenarthrans [51] and other slow arboreal therians considered here. Large vascular canals are however present in *P. cinereus*, especially in the endosteal region [65] We hence tentatively propose that the slightly higher humeral CC variability and the extremely wide femoral CC range of *P. cinereus* could be explained by a more variable extension of the observed primary vascularization in this species. In any case, the CC pattern of *P. cinereus* seems to reflect different factors than those of the other mammals investigated here.

### c. Relationship of low CC and bone remodeling with low metabolic rate

Several bone microstructural features and bone remodeling were linked to metabolic rates [67–69]. Low CC was qualitatively observed in humerus, femur, and other postcranial elements of ‘tree sloths’ [12], suggesting that a systemic phenomenon as low metabolism plays a role [12]. Indeed, ‘tree sloths’ exhibit a strikingly low metabolic rate for their body size [70,71]. Moreover, low thyroid activity was reported for *Choloepus* [72]. Thyroid hormones contribute to metabolism [73] and bone growth [74] regulation. Accordingly, intense but balanced remodeling through adulthood, responsible for low CC, was inferred for ‘tree sloths’ [12]. *Cy. didactylus* convergently appears to have acquired low CC. Beside evidence of slow arboreal ecology [54–56], *Cy. didactylus*’ metabolism was reported as lower than expected for its mass (but not as low as *Bradypus*; [56]). This suggests that slow arboreality/hypometabolism could be responsible for low CC in ‘tree sloths’ and *Cy. didactylus*.

Moderately low CC was independently acquired in palaeopropithecids (especially *Palaeopropithecus*) and *Megaladapis*. Despite the impossibility to directly measure metabolic activity in subfossil lemurs, several clues point toward hypometabolism [17]. Energy saving strategy is generally related to folivory [75], and this trophic adaptation was inferred for both palaeopropithecids and *Megaladapis* [17,19,76]. Moreover, tooth enamel accretion periodicity (Retzius periodicity) suggests possibly low metabolic activity [18,67,77]. We hypothesise that their hypometabolism is related to their actively remodeled cortex, which is in turn responsible for the relatively low CC in these taxa.

‘Lorisidae’ slow arboreal lifestyle is accompanied with low body mass-corrected metabolic activity [78]. It distinguishes them from other strepsirrhines, which are generally hypometabolic [79]. This condition mirrors that of ‘tree sloths’ and their distinctively low metabolism within the generally hypometabolic Xenarthra [71]. Nevertheless, no low CC was found in ‘Lorisidae’ and they do not show differences in CC to Galagidae. Both taxa follow the general therian condition. Differences from galagids in the pattern of intracortical remodeling were identified on osteo-histological sections and ascribed to the slow arboreal locomotion of ‘Lorisidae [60]. Positional behaviour could thus preponderantly explain cortical microstructure and intracortical remodeling in this clade, with metabolism considered a negligible factor [60].

CC in koala and wombats unlikely is explained in terms of lifestyle alone. *P. cinereus* belongs to a generally hypometabolic clade, the marsupials, and its unusually low metabolism stands out [64]. Although their terrestrial lifestyle is distinct from slow arboreal, wombats are known for their energy conserving adaptations [80]. We found that Vombatiformes’ CC deviates from the general mammalian pattern, with higher values in the humerus than in the femur in both taxa. To explain the relatively higher CC in the wombats’ humerus in respect to the femur, we propose that biomechanical demands for high CC related to their forelimb dominated scratch-digging, could locally act on the humerus [66]. The CC discrepancy between humeral and femoral mean CC remains unclarified in *P. cinereus*, especially when considering a systemic explanation. However, its inter-limb difference in mean CC is less extreme than in wombats, and the most striking aspect of koala’s CC is its variability. As shown by histology, *P. cinereus* is clearly distinct from other slow arboreal therians with extremely low bone remodeling activity [65]. We hypothesise that the CC pattern of *P. cinereus*, distinct from those of other slow arboreal mammals, does not reflect bone remodeling and metabolic factors.

## 5. Conclusions

Adult ‘tree sloths’ are characterised by low humeral and femoral CC. The trait was previously linked to highly active bone remodeling and low metabolism, related to their slow arboreal lifestyle. Extinct sloths sampled here show high levels of CC, thus low CC likely represents a recent convergence in ‘tree sloths’, reflecting their convergent lifestyle. Their extreme low CC are not paralleled by other slow arboreal/hypometabolic mammals. Nevertheless, relatively low CC identified in *Palaeopropithecus* and *Megaladapis* appears to be a convergent trait shared with ‘tree sloths’. This finding, together with the striking evidence of low CC in the silky anteater (*Cyclopes didactylus*), supports a driving role of slow arboreality/low metabolism on CC. However, the absence of a clear overall pattern in all analysed slow arboreal/hypometabolic therians calls for additional or alternative explanations. Future studies, possibly including a more diverse mammal sample and also non-arboreal hypometabolic mammal species, ideally complemented with experimental physiological evidence, are required to shed light on the functional significance of low CC.

## Data accessibility

The analysed dataset is available on Figshare Digital Repository (https://doi.org/10.6084/m9.figshare.12896249.v4) [48]. The Museum für Naturkunde (Berlin, Germany) stores raw CT data, making them available upon reasonable request.

## Authors’ contribution

F.A. collected the sample, acquired, processed and analysed µCT data, performed statistical analysis and drafted the manuscript. All authors designed the study, interpreted the data, and contributed to the writing and editing of the manuscript.

## Acknowledgements

We would like to thank curators and assistant curators who allowed visits to collections and specimens access: Frieder Mayer, Christiane Funk and Anna Rosemann (ZMB), Eva Bärmann (ZFMK), Frank Zachos and Alexander Bibl (NMW), Stefan Merker (SMNS), Anneke van Heteren (ZSM), Guillaume Billet (MNHN), Neil Duncan (AMNH), Sara Ketelsen (AMNH), Vanessa Rhue (YPM-PU), Daniel Brinkman (YPM-PU), Adam Ferguson (FMNH), William Simpson (FMNH), Matt Borths (DPC) and Catherine Riddle (DPC). Moreover, we acknowledge Kristin Mahlow and Martin Kirchner (Museum für Naturkunde, Berlin), Renaud Lebrun (MRI-ISEM, Montpellier), Justin Gladman (SMIF, Durham, NC, USA), April Isch Neander and Zhe-Xi Luo (University of Chicago, IL, USA) for allowing access to Micro-CT scanners, providing precious assistance. We acknowledge the MRI platform member of the national infrastructure France-BioImaging supported by the French National Research Agency (ANR-10-INBS-04, «Investments for the future»), the labex CEMEB (ANR-10-LABX-0004) and NUMEV (ANR-10-LABX-0020). This work was performed in part at the Duke University Shared Materials Instrumentation Facility (SMIF), a member of the North Carolina Research Triangle Nanotechnology Network (RTNN), which is supported by the National Science Foundation (Grant ECCS-1542015) as part of the National Nanotechnology Coordinated Infrastructure (NNCI).

